# Humans integrate gaze and decision cues for inferring preferences in social interactions

**DOI:** 10.64898/2026.07.09.737460

**Authors:** Mrugsen Gopnarayan, Sophie Bavard, Erik Stuchlý, Sebastian Gluth

**Affiliations:** Department of Psychology and Hamburg Center of Neural and Cognitive Systems, University of Hamburg, Von-Melle-Park 11, Hamburg, 20146, Germany; Paris Brain Institute - ICM, Hopital de la Pitie-Salpetriere, Paris, France

**Author notes:** Contributing authors.

**Keywords:** social cognition, decision-making, eye-tracking, Bayesian inference, value-based choice, bargaining, Game Theory

## Abstract

Social decision-making depends on inferring others’ hidden preferences from observable behavior. Yet it remains unclear how humans combine choices with process cues such as response times and gaze when learning about others in real-time interaction. Here we combine a novel multi-attribute bargaining task with eye-tracking and show that multiple decision-process cues support preference inference. Across 75 buyer-seller dyads, buyers’ acceptance rates tracked offer utility, rejection speed reflected decision confidence, and first fixations preferentially targeted the highest-weighted attribute. Sellers adapted subsequent offers using choices, response times, and, when available, gaze cues. A hierarchical inference and choice model suggested that sellers balanced expected utility with expected information gain and updated their beliefs in a Bayesian manner. Although gaze access did not improve overall performance, it changed how sellers used attentional information. These findings shed light on how humans infer others’ hidden preferences from decision dynamics in real-time social interaction.

## 1 Introduction

Humans are social learners by nature. We rely on behavioral signals to make sense of others’ internal states, including their intentions, beliefs, and preferences. Understanding the inferential mechanisms of human social inference is a longstanding challenge in cognitive science [1, 2]. While much is known about how people learn from the choices others make [3, 4], evidence indicates that decision dynamics, such as response times (RTs) and eye movements (gaze), can also be highly informative about the decision process [5–9]. In value-based decision-making, RTs have been found to correlate with preference strength, decision difficulty, and confidence. For example, slower decisions often accompany difficult or ambiguous choices, while fast decisions suggest more clear-cut preferences [10, 11]. Eye movements can also reflect preferences and predict choices: people tend to gaze longer at items they eventually choose, and, in multi-attribute settings, allocate attention to important attributes [12–16]. Furthermore, gaze not only reflects preferences but can also causally influence decisions, although meta-analytic evidence suggests that these effects are modest and depend on how attention is manipulated [17–19].

There is also evidence that humans can learn preferences from these decision-process cues, especially from RTs [5, 20], but also from gaze [21]. However, the mechanisms by which people use and combine these cues to make social inferences remain unclear. These additional process cues may play an even greater role in interactive social settings, where intentions and preferences can change over time and information from choices may be insufficient. Yet few studies have examined how dynamic cues are used in interactive, multi-round social contexts, where one must adapt behavior based on inferred latent states of another person. In such settings, individuals must make not only inferences but also predictions and counterfactual evaluations, often with incomplete feedback. A prime example is bargaining, where a seller must learn what a buyer wants through their reactions to offers. Sellers receive limited explicit feedback, such as accept/reject decisions, but potentially richer implicit signals via RT and gaze. The critical question is therefore whether and how humans exploit these cues to improve interactive learning.

In this study, we asked whether real-time access to others’ eye movements, alongside their choices and decision speed, can facilitate preference inference in social interactions. We designed a novel multi-attribute bargaining task (Figure 1), a multi-round buyer-seller setting with gaze access as a between-group manipulation. Sellers had to infer the buyer’s unknown attribute weights from offer feedback and follow up with better offers. In the Gaze group, sellers viewed a live overlay of the buyer’s gaze, enabling them to see which attribute the buyer was currently looking at. In the No Gaze group, sellers observed only the buyer’s decisions and the timing of those decisions.

**Fig. 1.**
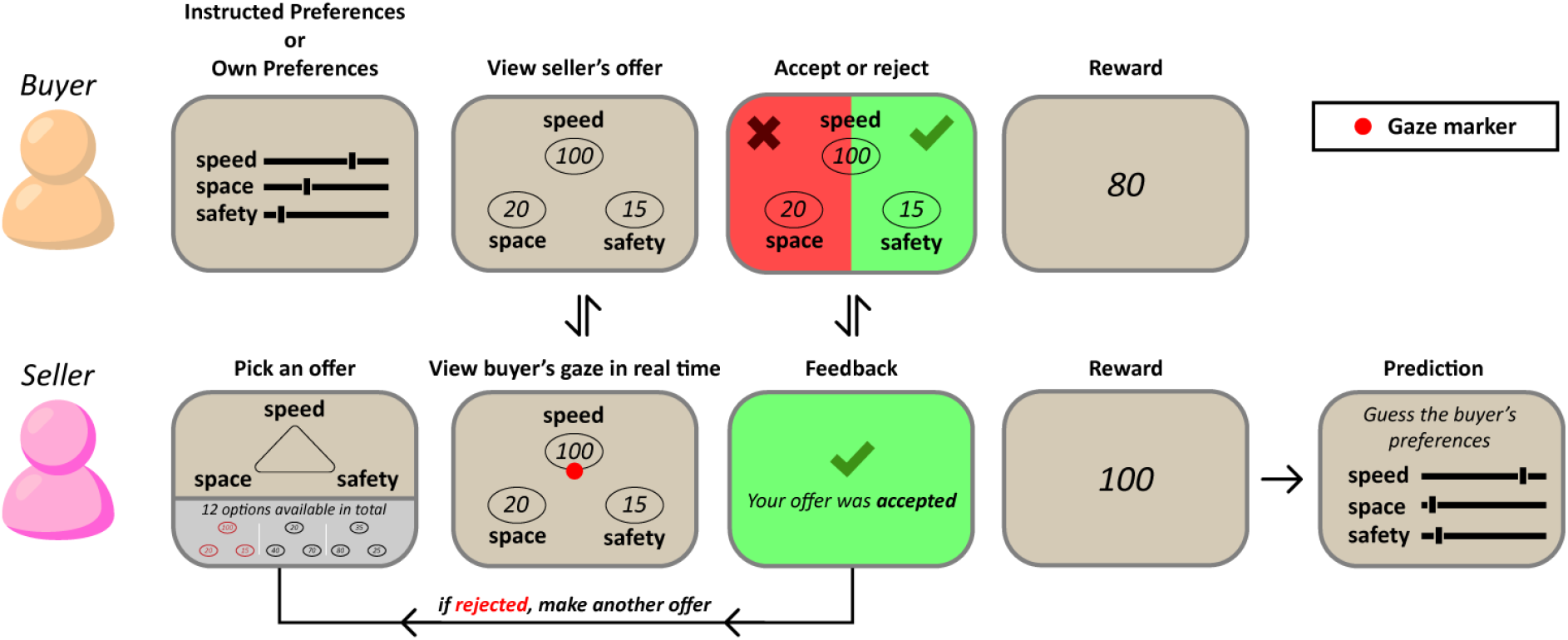
Schematic Illustration of the Multi-Attribute Bargaining Task. Two players interact: a Buyer (B) with latent attribute weights and a Seller (S) who must infer them. Panels depict progression of a single trial for item category ‘Car’. B’s attribute weights are determined either by instructed weights or by their own natural weights (here, B mostly cares about the speed of the car). S chooses an offer from a fixed menu of 12 options that vary only in how the scores are distributed among the three attributes. B views the proposed option. In the Gaze group, S sees B’s gaze in real time as a marker over the three attributes; in the No Gaze group this marker is absent. B accepts or rejects; the decision is communicated immediately, revealing reaction time to S without an explicit timer. Reward presentation follows: B earns the utility of any accepted offer (attributes scaled 0-100) or a baseline of 50 if all offers are rejected; S’s payoffs are computed separately, based on whether the product was sold or not. S reports their beliefs about B’s attribute weights. The sequence repeats across 20 categories, with a unique buyer weight vector in each category.

To elucidate the underlying choice strategies and cognitive mechanisms, we developed computational models for both agents. Buyer choices and RTs were modeled with a drift diffusion model (DDM), a sequential-sampling model in which noisy evidence accumulates over time until a decision boundary is reached. Sellers were modeled as Bayesian learners who updated beliefs about the buyer’s attribute weights from observed outcomes. The model asked whether sellers simply pursued high-utility offers or also selected offers that would be informative about the buyer’s hidden weights. It also quantified how strongly gaze patterns were treated as evidence about attribute weights.

We tested the following pre-registered hypotheses: (1) buyers’ attribute weights are reflected not only in their choices but also in their RTs and eye movements; (2) sellers exploit all of these cues to make more informed offers; and, consequently, (3) sellers in the Gaze group outperform those in the No Gaze group. Our findings were largely consistent with the first two predictions. Gaze and RTs reflected buyer weights, and sellers used these cues to adapt their offers. Surprisingly, while gaze access affected sellers’ offers, it did not lead to overall higher earnings, suggesting that real-time gaze streams impose substantial cognitive demands that offset their informational benefits when choices and RTs already provide rich feedback. Model-based analyses revealed nuanced seller strategies involving both exploitation and active inference, offering a richer picture of how people learn from decision-process information in social contexts.

## 2 Results

We analyzed behavior and model fits from 150 participants (75 buyer-seller pairs) who completed a multi-attribute bargaining task (Figure 1). Pairs interacted across 20 product categories (e.g., cars, laptops), each defined by three attributes (e.g., speed, comfort, safety). In each category, the seller could sequentially propose up to four offers from a fixed menu of twelve options. Buyer utilities were defined as the attribute-weighted sum (scaled 0-100), and buyers earned a reward proportional to the accepted offer’s utility or a baseline of 50 points if they rejected all four offers. Sellers observed decisions in real time, so that RTs were implicitly signaled by the interval between offer and response. In the Gaze group (38 pairs), sellers also viewed a live fixation overlay on their screen; in the No Gaze group (37 pairs), only decisions and timing were visible.

Buyers’ mean acceptance utility was 60.7 (SD = 7.6), significantly above the 50-point baseline (*t*(2399) = 68.99, *p <* .001), indicating utility-driven decision-making. For accepted offers, sellers earned a mean of 86.9 points (SD = 14.3), significantly above chance (*t*(2399) = 126.66, *p <* .001).

### 2.1 Choice and Response Times

Buyer decisions were systematically linked to offer utility. A logistic regression revealed that acceptance probability increased with utility (slope *B* = 0.154, SE = 0.007, *z* = 23.75, *p <* .001; inflection point at utility = 60.0; Fig. 2a). The model explained substantial variance (pseudo-*R*^2^ = 0.22). Across the four offer iterations, there was a modest downward shift in the acceptance threshold (mean shift *≈* 3 utility points per iteration), consistent with buyers strategically relaxing criteria as the offer limit approached.

**Fig. 2.**
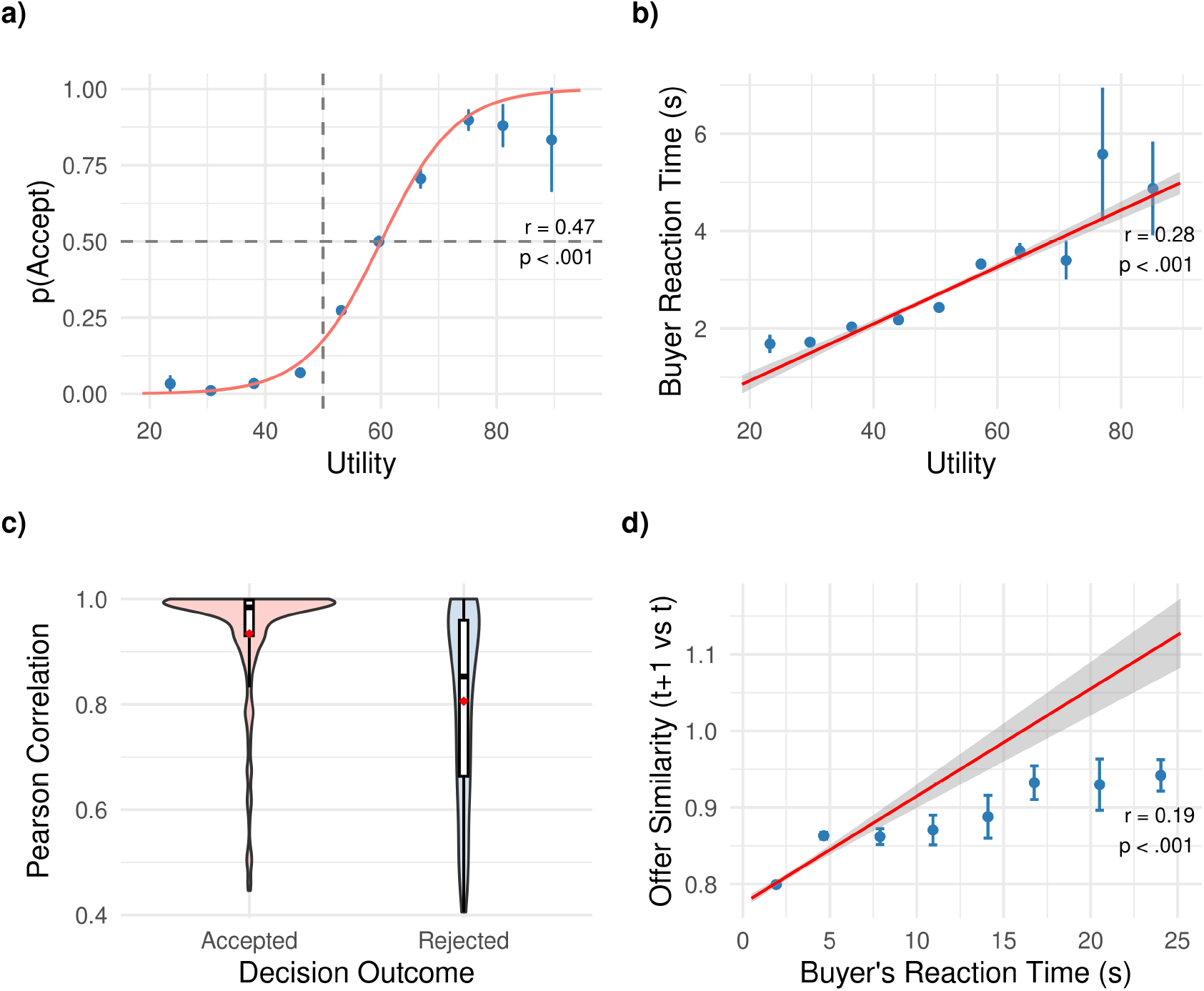
Choice and response time behavior. **a**, Buyer acceptance probability increases with offer utility. Colored curves show fitted logistic functions for each of the four offer attempts within a category. **b**, Rejection RTs increase with utility, with low-utility offers rejected more quickly than borderline offers. **c**, Sellers learn from choice outcomes: Pearson correlations between final offers and sellers’ preference estimates are higher when offers are accepted versus rejected. **d**, Sellers adapt to RT signals: faster rejections are associated with lower cosine similarity between the current offer and the next offer, indicating larger offer changes.

RTs provided additional information about buyers’ decision confidence. For rejected offers, RTs increased with utility (linear mixed-effects model: *B* = 0.061, SE = 0.005, *t* = 12.51, *p <* .001; Fig. 2b), indicating that clearly low-utility offers (utility *<* 40) were rejected quickly (mean RT = 1.8 s), whereas borderline offers (utility 50-60) elicited slower rejections (mean RT = 2.5 s).

Turning to the behaviour of sellers, we find evidence that they learned from the buyer’s behaviour described above. Sellers learned from binary choice outcomes. When the final offer in a category was accepted, the Pearson correlation between the final offer and the seller’s preference estimate was substantially higher (M = 0.79, SD = 0.44) than when all offers were rejected (M = -0.03, SD = 0.72; *t*(880.87) = 21.16, *p <* .001, Cohen’s *d* = 1.33; Fig. 2c). Sellers also adapted to RT signals: the buyer’s rejection speed in trial *t* predicted the similarity between the current offer and the seller’s next offer in trial *t* + 1 (*B* = 2.47, SE = 0.22, *t* = 11.33, *p <* .001; Fig. 2d), demonstrating that sellers correctly interpreted fast rejections as signaling low utility.

### 2.2 Gaze Patterns Reveal Attribute Weights

Eye-tracking data (N = 7,157 trials) revealed systematic relationships between gaze and attribute weights. Buyers first fixated their highest-weighted attribute in 58% of trials, significantly above chance (33%; *t*(7156) = 42.28, *p <* .001, 95% CI [0.57, 0.59]; Fig. 3a). In contrast, first fixations landed on the second-highest attribute in 25% of trials (*t*(7156) = -16.62, *p <* .001) and the lowest-weighted attribute in 17% of trials (*t*(7156) = -36.29, *p <* .001), both significantly below chance. Last fixations were less diagnostic and showed no consistent weight alignment.

**Fig. 3.**
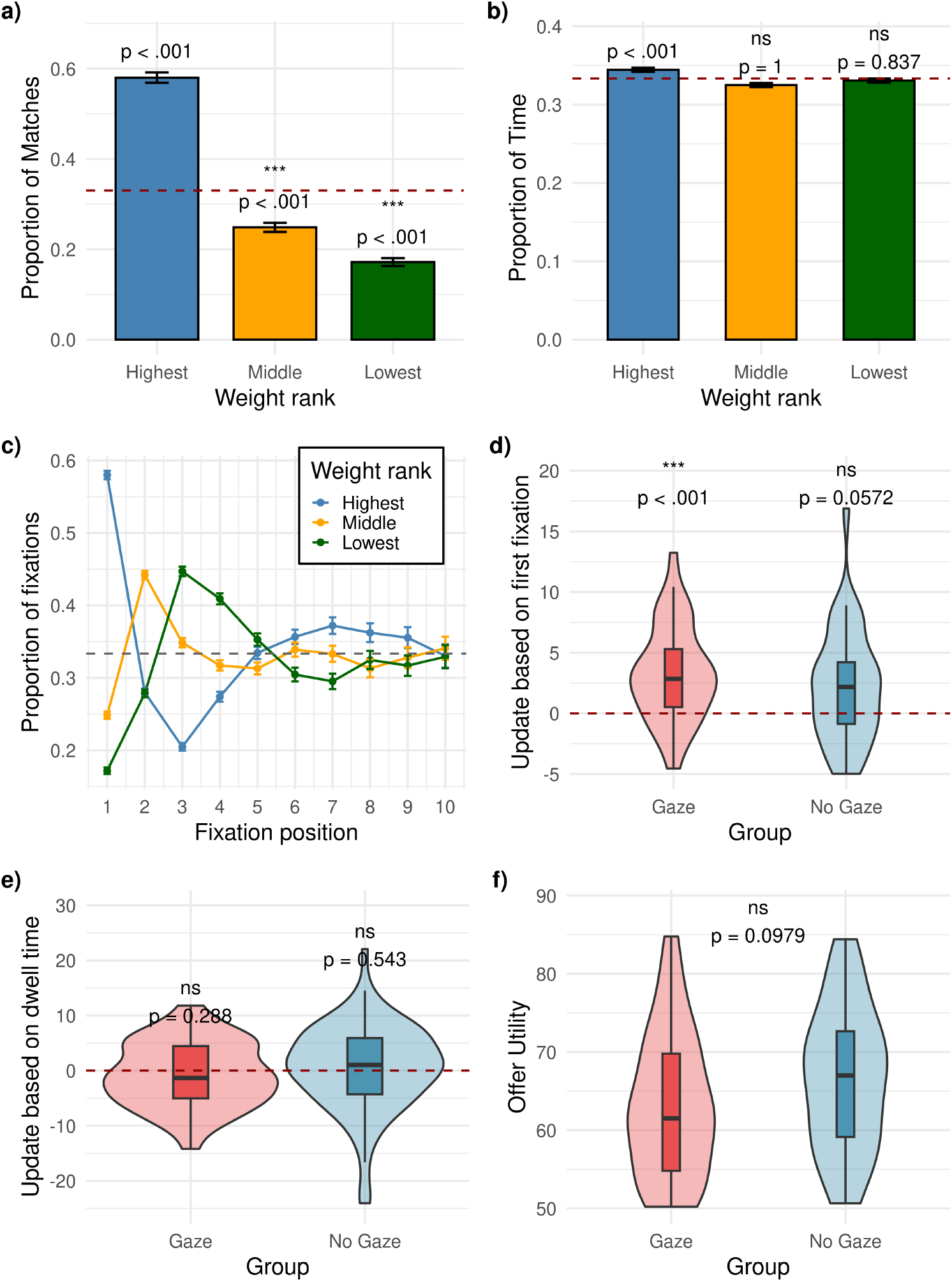
Gaze patterns and seller adaptation. **a**, First fixations landed mostly on the highest-weighted attributes (58% vs. 33% chance), with middle-weighted (25%) and lowest-weighted (17%) attributes fixated less often than chance (dotted red lines). **b**, Dwell time shows modest differentiation by attribute weight, with highest-weighted attributes receiving 34% of viewing time. **c**, Fixation order profiles reveal early attentional bias that diminishes across fixation sequence. **d**, Sellers in the Gaze group increased the score of first-fixated attributes in subsequent offers. **e**, Dwell time did not affect sellers’ updates. **f**, Gaze access did not increase overall earnings (Gaze: M = 60.1; No Gaze: M = 63.8).

Dwell time (total fixation duration per attribute) showed modest differentiation by attribute-weight rank (Fig. 3b). Highest-weighted attributes received a mean of 34% of total viewing time compared to chance level of 33% (*t*(7156) = 4.26, *p <* .001, 95% CI [0.34, 0.35]), but this effect was weak. Fixation order was more informative (Fig. 3c): the probability of fixating the highest-weighted attribute declined across fixation sequence (1st fixation: 58%, 2nd: 38%, 3rd: 29%), suggesting that early attention reliably indexed weights [16].

Sellers in the Gaze group exploited first-fixation information strategically. After unsuccessful offers, sellers increased the score of the first-fixated attribute in their next offer by an average of 3.0 points (SD = 4.1). This adaptation was significantly greater than zero (*t*(37) = 4.52, *p <* .001; Fig. 3d), whereas updates based on dwell time were not significant (mean change = -0.7 points, *t*(75) = -1.07, *p* = .288; Fig. 3e).

Despite this strategic use of gaze, overall performance did not differ between groups (Fig. 3f). Sellers in the Gaze group earned a mean of 60.1 points per negotiation (SD = 12.7), compared to 63.8 points (SD = 11.9) in the No Gaze group, a non-significant difference (*t*(72.88) = -1.31, *p* = .20, Cohen’s *d* = -0.30).

### 2.3 Natural Attribute Weights Facilitate Better Performance

Performance systematically differed between instructed and natural-weight blocks. In blocks where buyers used their own natural attribute weights, offer utility was higher (M = 55.7, SD = 1.9) than in blocks with instructed weights (M = 52.0, SD = 1.5; paired *t*(74) = 13.81, *p <* .001, Cohen’s *d*_*z*_ = 1.59; Fig. 4b). This advantage likely indicates that sellers could use knowledge of their own attribute weights to anticipate buyers’ weights, consistent with evidence that prior preferences can shape social learning [22]. Sellers’ offers improved in the within each category (Fig. 4c). for both Gaze and no Gaze group in the instructed block, demonstrating robust within-category learning from binary feedback and RTs alone.

**Fig. 4.**
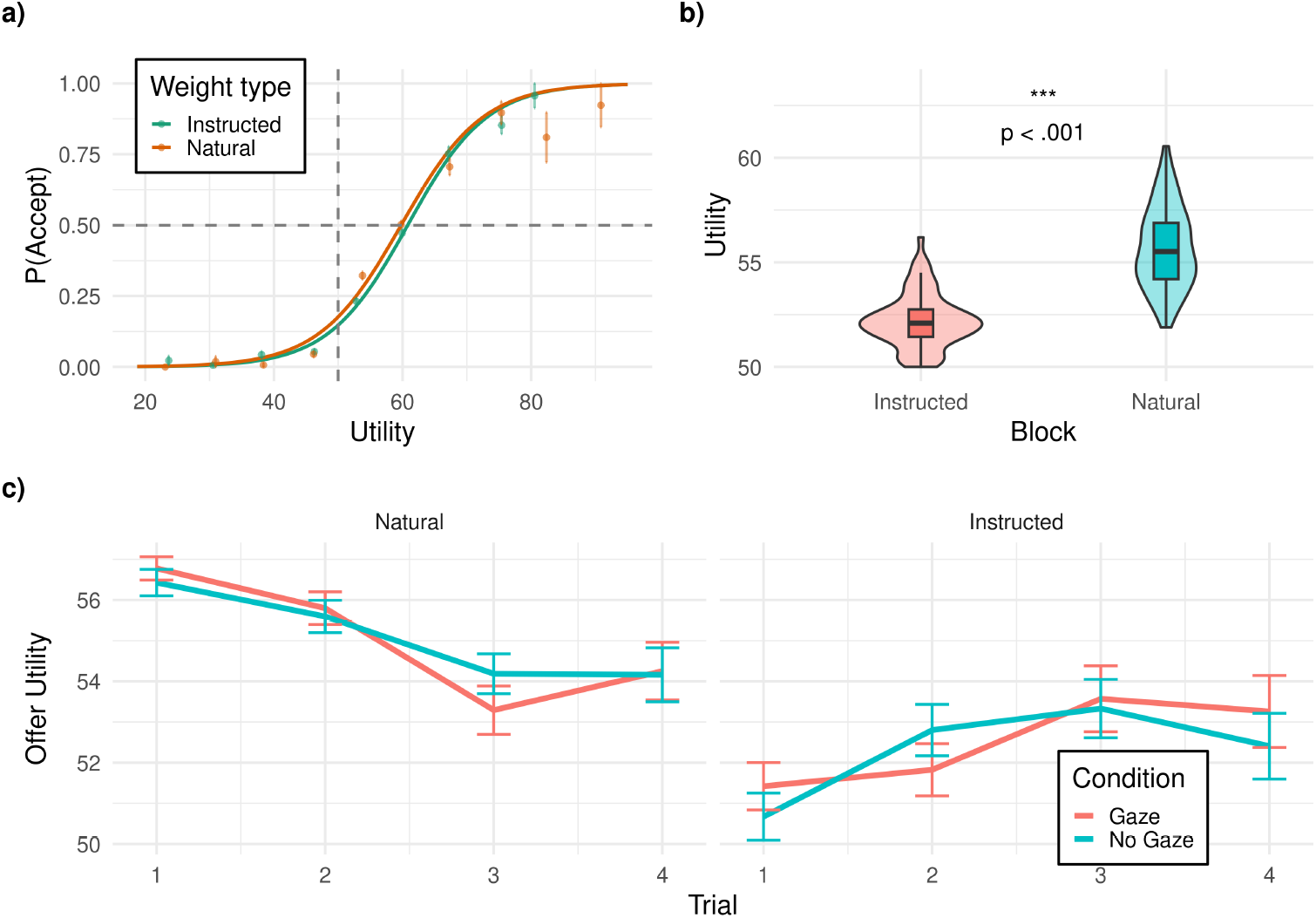
Natural attribute weights facilitate better performance. **a**, Buyer acceptance curves are similar across weight types. **b**, Offer utility is higher in natural-weight blocks than in instructed-weight blocks. **c**, Offer utility improves across the four offer attempts within each category for the instructed block but not for the natural block

Buyer acceptance curves were similar across block types (Fig. 4a). The logistic utility slope was 0.157 (SE = 0.007) for natural-weight blocks and 0.162 (SE = 0.006) for instructed-weight blocks, with no reliable interaction between utility and block type (*B* = -0.005, SE = 0.010, *z* = -0.52, *p* = .603).

### 2.4 Computational Models Reveal Information-Utility Trade-off

We formalized seller inference using a Bayesian social inference model. Sellers maintained a posterior belief distribution over buyer attribute weights and selected offers that maximized a weighted combination of expected utility and expected information gain (IG):

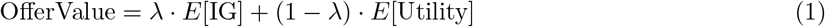

where *λ* ∈ [0, 1] is the Information-Utility Tradeoff parameter (*λ* = 1: pure information seeking; *λ* = 0: pure utility maximization). Information gain quantifies how much an offer is expected to reduce uncertainty about the buyer’s attribute weights. The term is expected because sellers evaluate each offer before observing the buyer’s response, averaging over possible responses under their current belief state (see Methods). Hierarchical Bayesian inference recovered meaningful parameter estimates from the two-stage model (Fig. 5). In the first stage, buyer-level parameters were estimated from buyers’ choices, RTs, and gaze sequences. These buyer parameters were then passed forward to the seller model, which estimated how sellers traded off utility and information and how strongly they used gaze. The buyer-level utility cutoff *µ* was 61.67 in the No Gaze group (95% CI [59.97, 63.44]) and 63.21 in the Gaze group (95% CI [61.91, 64.51]). Boundary separation was similar across groups (No Gaze: *a* = 3.60, 95% CI [3.32, 3.87]; Gaze: *a* = 3.76, 95% CI [3.47, 4.05]), as were drift scaling (No Gaze: *v* = 0.047, 95% CI [0.041, 0.053]; Gaze: *v* = 0.050, 95% CI [0.044, 0.057]) and gaze sensitivity in the buyer gaze model (No Gaze: *α* = 2.27, 95% CI [1.74, 2.67]; Gaze: *α* = 2.18, 95% CI [1.67, 2.60]). Altogether, buyers’ decision strategy did not significantly differ between Gaze and No Gaze groups.

**Fig. 5.**
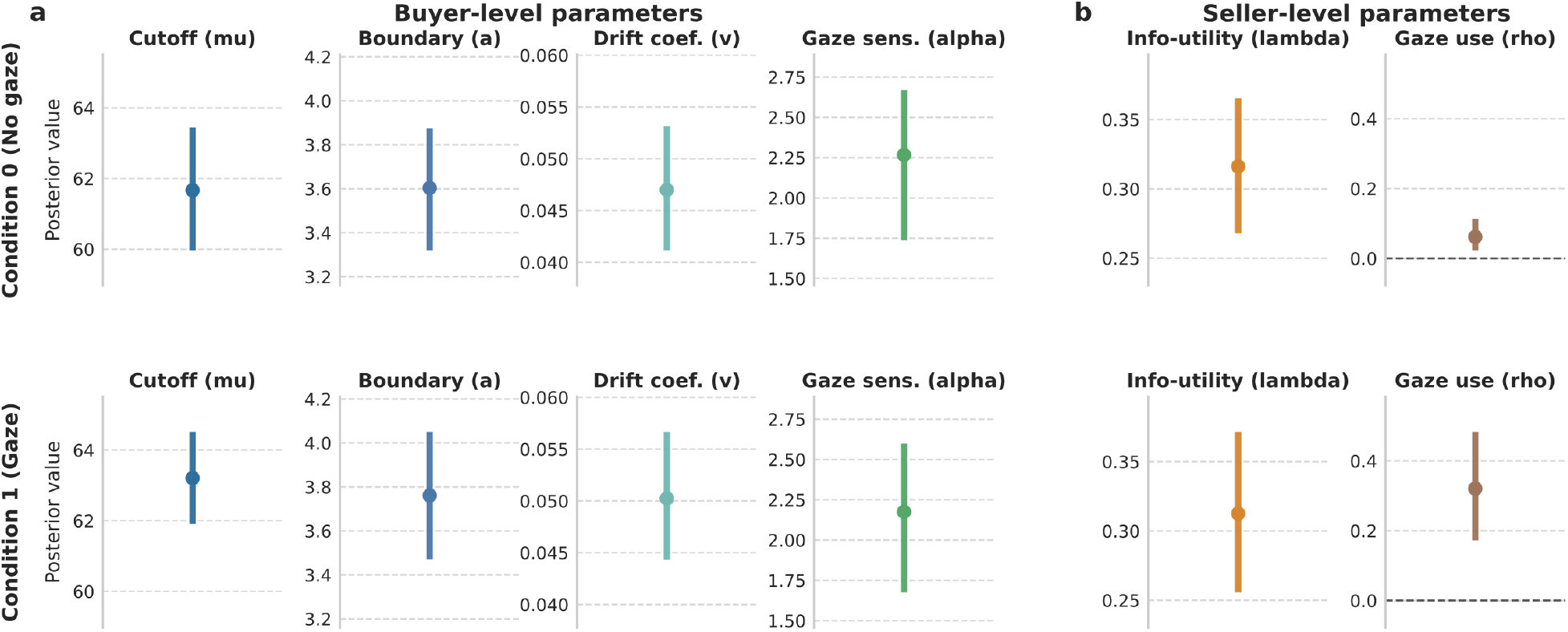
Two-stage computational model parameter estimates. **a**, Buyer-level parameters estimated from buyers’ choices, RTs, and gaze sequences: utility cutoff *µ*, boundary separation *a*, drift coefficient *v*, and gaze sensitivity *α*. **b**, Seller-level parameters estimated after passing the buyer-level parameters forward to the seller model: the Information-Utility Tradeoff parameter *λ* and gaze-use parameter *ρ*. Points show posterior means and vertical intervals show 95% credible intervals.

At the seller level, the Information-Utility Tradeoff parameter *λ* was reliably greater than zero in both groups (No Gaze: *λ* = 0.316, 95% CI [0.268, 0.365]; Gaze: *λ* = 0.313, 95% CI [0.256, 0.371]), indicating that sellers were information seeking rather than purely reward-maximizing. The gaze-use parameter *ρ* captured how strongly the gaze likelihood was weighted during seller inference. It was close to zero in the No Gaze group (*ρ* = 0.062, 95% CI [0.023, 0.113]) and larger in the Gaze group (*ρ* = 0.321, 95% CI [0.172, 0.483]), consistent with sellers incorporating visible fixation patterns into their inferences when gaze was available.

Taken together, these results suggest that sellers made use of all three available cues (choices, RTs, and gaze) to infer buyers’ attribute weights in an interactive, strategic environment. They integrated information and updated their beliefs in a Bayesian way.

## 3 Discussion

We asked whether decision-process signals – choices, response times, and eye movements – carry exploitable information about latent preferences in an interactive exchange. Our findings provide a clear answer: all three streams encoded preference-relevant information, and human observers spontaneously used them to adapt their behavior. Buyers’ acceptance probability increased sigmoidally with offer utility, their rejection speed scaled inversely with utility (fast rejections signaled clear-cut low utility), and their first fixations reliably targeted highest-weighted attributes. Sellers exploited these signals: they changed offers more strongly after fast rejections and, when given access to gaze, tilted subsequent offers toward first-fixated attributes. Yet despite this strategic use of attention, gaze access did not translate into higher overall earnings. This pattern suggests that while process data are informative in principle, real-time attentional streams can impose complexity that offsets their benefits when choices and response times already provide rich feedback.

Computational modeling revealed that sellers valued information, not only utility. The Information-Utility Tradeoff parameter *λ* was reliably greater than zero in both groups, indicating that sellers asked not only “which offer maximizes expected utility?” but also “which offer would I learn the most from?” This shows they engaged in counterfactual evaluation. This information-seeking behavior is consistent with active inference frameworks [23, 24] in which agents select actions partly to reduce uncertainty about hidden states. The gaze-use parameter *ρ* was larger in the Gaze group, confirming that sellers incorporated visible fixation patterns into their weight inferences. Together, these modeling results suggest that sellers maintained probabilistic representations of buyer weights and weighted information gain alongside immediate utility.

These findings extend several lines of prior work. Studies of RT-based inference have shown that observers can infer hidden preferences from decision speed, sometimes even when choices are unavailable [5, 20]. Our results extend this logic to an interactive setting in which observers must use decision speed in real-time to adapt their own future actions. Likewise, work on attention and value-based choice shows that gaze reflects and can shape valuation [6, 17, 19]; here, gaze served as a social cue that another person could exploit. Finally, the finding that first fixations were more diagnostic than dwell time connects the present task to models of multi-attribute search, in which attention is allocated in a goal-directed manner to information that is currently useful for choice [16].

However, the attribute space comprised only three dimensions and a discrete menu of twelve offers; real negotiations often involve continuous features, budgets, and explicit communication. Our participants could not verbally negotiate, which was essential to avoid the overshadowing of non-verbal cues, but it removes a major channel through which preferences are typically revealed. The four-offer horizon provided limited scope for long-term exploration to pay off, potentially underestimating the value of information-seeking strategies in extended interactions. Despite these constraints, the fundamental insight – that humans integrate multiple process cues, update beliefs in a Bayesian-like fashion, and also make decisions specifically to inform their beliefs about others – likely generalizes beyond our specific paradigm.

These findings have broader implications for understanding social cognition and for designing human-AI interaction systems. Our results demonstrate that humans possess sophisticated mechanisms for inferring preferences from decision dynamics, extending the growing literature on “mind-reading” from choice data alone [2] to encompass temporal and attentional signatures. From an applied perspective, the observation that first-fixation cues are more actionable suggests design principles for attention-aware interfaces: distilling gaze into simple, early indicators useful for preference inference. Future work should examine how well current AI systems, such as large language models, perform at making such inferences. Training such systems to infer latent states from decision process data [25] points toward human-AI interaction systems in which AI interprets complex signals (such as gaze or neural activity). Such systems could enhance negotiation, personalized recommendation, and clinical assessment of social cognition in conditions where preference inference is impaired.

## 4 Methods

### 4.1 Participants

A total of 158 participants took part in the study (79 pairs; mean age = 24.9 years, SD = 6.9; 113 female). Out of which 4 pairs were excluded from the analysis based on pre-registered criterion on performance and data quality. All participants were naive to the purpose of the study and gave written informed consent. Ethical approval was obtained from the Local Ethics Committee of the Faculty of Psychology and Human Movement Sciences at the University of Hamburg (approval number 2024 010). Each participant received a fixed show-up fee of 26 Euros or 2 course credit points plus a performance-based bonus (max 6 Euros) proportional to their total reward in the task.

### 4.2 Task Procedure

Participants completed a multi-attribute bargaining task. The task comprised 20 product categories (e.g., cars, laptops, smartphones), each defined by three unique attributes (e.g., for cars: speed, comfort, safety). Attribute labels and their mapping to categories were counterbalanced across participants. For each category, 12 product options were generated with unique attribute scores, randomly distributed from 0-100, with higher scores indicating better attribute-specific scores of the offer.

Before each trial, the buyer viewed their weights for the three attributes. In instructed blocks, these weights were randomly assigned from a set of predetermined triplets (e.g., 60%-30%-10%, 50%-30%-20%); in natural blocks, weights reflected the buyer’s own ratings, which they reported via three sliders. The seller, unaware of the buyer’s weights (but aware of the block type, instructed vs. natural), selected one of the 12 options to propose. The buyer then viewed the offer and decided to accept or reject it via key-press (‘right arrow’ = accept, ‘left arrow’ = reject). RTs were recorded from offer onset to key-press.

If the buyer rejected the offer, the seller could propose up to three additional offers (maximum four per category). If any offer was accepted, the category round ended and both players received feedback on their earnings. If all four offers were rejected, the buyer received a fixed baseline of 50 points, and the seller received 0 points. For all 12 offers we calculated their utility (computed as *u* = *w*_1_ ·*a*_1_ + *w*_2_ ·*a*_2_ + *w*_3_ ·*a*_3_, where *w*_*i*_ are the weights and *a*_*i*_ are the attribute scores). For accepted offers, the buyer earned a reward from 0-100 based on the utility based ranking of the offers and the seller’s reward was proportional to the buyer’s reward (computed as 0.8 * *BuyerReward* + 20). This payoff structure incentivized sellers to maximize buyer utility while encouraging buyers to only accept offers exceeding the baseline.

Between each seller decision, a fixation cross appeared for 1.5 *±* 0.5 s. In the Gaze group, while the buyer viewed the offer, a red cursor on the seller’s screen updated in real time to indicate the buyer’s current gaze position. In the No Gaze group, sellers saw only a static offer display during the buyer’s decision phase.

### 4.3 Experimental Setup

Participants were randomly assigned to the role of buyer or seller and were assigned to one of two groups: Gaze group (seller has gaze access) and No Gaze group (no gaze access). They were placed in different rooms and did not see each other before the task, preserving anonymity. Before the task began, participants followed a plastic hand-waving protocol in which they waved a set of plastic hands to each other. This procedure made the presence of another participant salient without revealing either participant’s identity.

### 4.4 Experimental Design

Each pair was randomized into one of two groups. In the Gaze group, sellers saw a live display of the buyer’s gaze. In the No Gaze group, buyers’ gaze was recorded but sellers observed only buyer decisions and timing, with no access to gaze data. Each session was divided into two blocks that manipulated how buyer attribute weights were determined. In instructed blocks, buyers were given an explicit set of weights for each attribute. In natural blocks, buyer weights reflected the buyer’s own ratings elicited via sliders for each attribute. The order of instructed and natural blocks was counterbalanced across participants.

### 4.5 Gaze Manipulation and Eye-Tracking

Buyers’ eye movements were recorded using an EyeLink 1000 Plus eye tracker (SR Research) with a sampling rate of 1000 Hz. In the Gaze group, sellers saw a red dot on their screen indicating the buyer’s current gaze position. This information was live-streamed, hence the eye-tracking data wasn’t pre-processed.

### 4.6 Pre-registration

This study was pre-registered on the Open Science Framework (OSF) under the title ‘Unravelling preferences of others through their eye movements in a Bargaining task’ (Gopnarayan, M., Bavard, S., Stuchlý, E., & Gluth, S., 2024). Link :(https://osf.io/x3jvc/overview).

### 4.7 Computational Modeling

#### Buyer Model (gaze-DDM)

We modeled buyer accept-reject decisions and RTs with a drift diffusion model (DDM) in which the drift rate depends on the utility of the offered option given the buyer’s latent attribute weights. For a given buyer and offer *o* with attribute vector **x**_*o*_, utility is

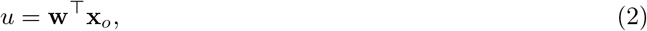

where **w** is a three-dimensional weight vector on the attribute space. This utility is mapped to a drift rate

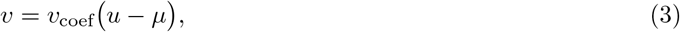

where *µ* is a cutoff point on the utility scale and *v*_coef_ scales utilities into drift rates. Evidence then accumulates toward an upper (accept) or lower (reject) boundary with separation *a*. The joint distribution of the binary decision *d* and response time *τ* is given by the DDM first-passage time density *f*_DDM_

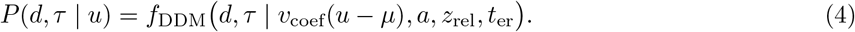

In the Gaze group, we additionally modeled the sequence of fixations within each trial. For a given trial and candidate weight vector **w**, the probability that fixation *ℓ* falls on attribute *i* was assumed to increase with the corresponding weight *w*_*i*_. Specifically, we used a softmax function over attributes with time-varying gaze sensitivity:

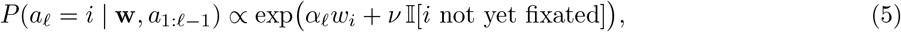

where *a*_*ℓ*_ is the attribute fixated at position *ℓ, α*_*ℓ*_ = *α/*(1 + *β*_*α*_(*ℓ* − 1)) controls how strongly early fixations align with the latent weights, and *ν* is a novelty bonus for attributes that have not yet been inspected on that trial. Positive values of *α* indicate that gaze is biased toward more strongly weighted attributes, whereas *α* close to zero implies that fixation sequences are largely independent of the underlying weights. In the two-stage modeling approach, buyer-DDM parameters (*µ, v*_coef_, *a, z*_rel_, *t*_er_) and gaze parameters (*α, ν*) were estimated first from buyer data and then passed forward to the seller model described below.

#### Seller Model (gaze-DDM inference)

We formalized seller inference as Bayesian learning over the three-dimensional buyer weight vector **w** = (*w*_1_, *w*_2_, *w*_3_), subject to Σ_*i*_ *w*_*i*_ = 1 and *w*_*i*_ ≥ 0. The seller represented uncertainty over **w** on a discrete grid of candidate weight vectors 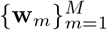 and maintained a posterior belief distribution *p*_*t*_(*m*) = *P* (**w** = **w**_*m*_ | data up to *t*) on this grid, initialized from a uniform prior.

On each trial, the seller observed: (i) their own offer choice, (ii) the buyer’s accept-reject decision and RT, and, in the Gaze group, (iii) the buyer’s fixation sequence. For a given grid point *m*, these observations define a joint likelihood

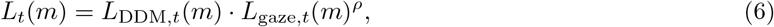

where *L*_DDM,*t*_(*m*) is the DDM likelihood of the buyer’s decision and RT under **w**_*m*_ (using the buyer-stage DDM parameters), *L*_gaze,*t*_(*m*) is the likelihood of the fixation sequence under the buyer-stage gaze model, and *ρ* controls how strongly sellers weight gaze evidence. When *ρ* = 0, gaze contributes no information to the seller’s update; larger values give the gaze likelihood more influence. We implemented sellers’ belief updates using the standard Bayesian update

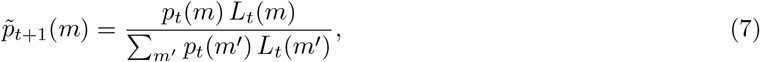

For offer selection, the seller evaluated each feasible option *o* using a combination of expected utility and expected information gain. Expected utility was defined as the posterior mean utility under the current belief

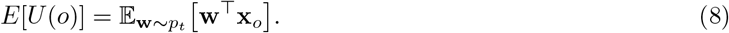

To quantify the informativeness of an offer, we used the DDM to approximate how much a rejection of that offer would sharpen the seller’s posterior. For each grid point *m* we computed the DDM-based rejection probability for offer *o, P*_rej_(*m, o*), and the implied posterior after a rejection

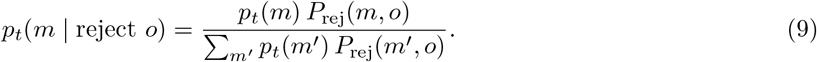

We then defined an information gain term as the reduction in entropy associated with such a rejection:

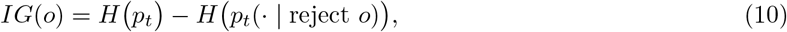

where *H*(*p*) = − *Σ*_*m*_ *p*(*m*) log *p*(*m*) is the Shannon entropy. Essentially, IG quantifies the expected change in belief about the attribute weights if an offer gets rejected (i.e., whether a rejection will inform the seller more or less about the buyer’s hidden attribute weights).

Finally, the seller traded off expected utility and information gain when deciding which option to propose:

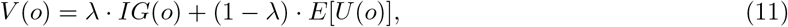

where *λ* ∈ [0, 1] is the Information-Utility Tradeoff parameter (*λ* = 0 corresponds to purely reward-maximizing behavior, *λ* = 1 to purely information-seeking behavior). Offers were selected via a softmax choice rule

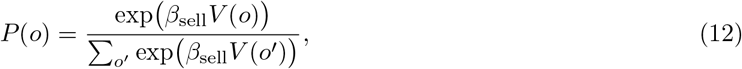

with inverse temperature *β*_sell_ controlling choice stochasticity. The DDM parameters (*µ, v*_coef_, *a, z*_rel_, *t*_er_) and gaze parameters (*α, ν*) used in the seller likelihood were fixed to the posterior estimates from the buyer-stage fits.

#### Hierarchical Bayesian estimation

We used a two-stage hierarchical Bayesian procedure in NumPyro. First, buyer-level parameters governing choices, RTs, and gaze sequences were estimated from buyer data. Second, these buyer-stage estimates were passed forward to the seller model, where the free seller-level parameters were the Information-Utility Tradeoff *λ* and the gaze-use parameter *ρ*. Both parameters were bounded in [0, 1] and modeled on the logit scale with weakly informative Normal hyperpriors, allowing participant-level variation around group-level means.

Posterior inference used the No-U-Turn Sampler (NUTS) with 8 chains, 500 warmup iterations, and 1,000 sampling iterations per chain. Convergence was assessed via 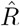 (all *<* 1.01) and effective sample size, and model fit was evaluated with posterior predictive checks on choice patterns, response times, and gaze adaptation across trials.

### 4.8 Statistical Analysis

All statistical analyses were conducted in R (version 4.3.0). For buyer choice data, we used generalized linear mixed models (GLMMs) with a logit link, including random intercepts for participants. For reaction time analyses, we used linear mixed-effects models (LMMs) with random slopes and intercepts for participants, implemented in the lme4 package. Models were fit using maximum likelihood, and significance was assessed via likelihood ratio tests comparing full models to null models (without the predictor of interest).

For eye-tracking analyses, we excluded trials with missing or invalid fixations (*<* 5% of data). First fixation was defined as the first gaze landing on any of the three attribute regions. Dwell time was computed as the sum of all fixation durations within each attribute region during the offer viewing period. Comparisons of first-fixation proportions to chance (33%) used one-sample *t*-tests.

Group comparisons (Gaze vs. No Gaze, natural vs. instructed) used independent-samples or paired *t*-tests as appropriate, with random effects for participants when data were hierarchically structured. Effect sizes were reported as Cohen’s *d* for mean differences and as *R*^2^ or pseudo-*R*^2^ for regression models. For all tests, *α* = .05 (two-tailed unless otherwise specified). All reported *p*-values are exact unless noted as *<* .001.

